# Reproducible genomics analysis pipelines with GNU Guix

**DOI:** 10.1101/298653

**Authors:** Ricardo Wurmus, Bora Uyar, Brendan Osberg, Vedran Franke, Alexander Gosdschan, Katarzyna Wreczycka, Jonathan Ronen, Altuna Akalin

## Abstract

In bioinformatics, as well as other computationally-intensive research fields, there is a need for workflows that can reliably produce consistent output, independent of the software environment or configuration settings of the machine on which they are executed. Indeed, this is essential for controlled comparison between different observations or for the wider dissemination of workflows. Providing this type of reproducibility, however, is often complicated by the need to accommodate the myriad dependencies included in a larger body of software, each of which generally come in various versions. Moreover, in many fields (bioinformatics being a prime example), these versions are subject to continual change due to rapidly evolving technologies, further complicating problems related to reproducibility. Here, we propose a principled approach for building analysis pipelines and managing their dependencies. As a case study to demonstrate the utility of our approach, we present a set of highly reproducible pipelines for the analysis of RNA-seq, ChIP-seq, Bisulfite-seq, and single-cell RNA-seq. All pipelines process raw experimental data, and generate reports containing publication-ready plots and figures, with interactive report elements and standard observables. Users may install these highly reproducible packages and apply them to their own datasets without any special computational expertise beyond the use of the command line. We hope such a toolkit will provide immediate benefit to laboratory workers wishing to process their own data sets or bioinformaticians seeking to automate all, or parts of, their analyses. In the long term, we hope our approach to reproducibility will serve as a blueprint for reproducible workflows in other areas. Our pipelines, along with their corresponding documentation and sample reports, are available at http://bioinformatics.mdc-berlin.de/pigx

## Introduction

Reproducibility of scientific workflows is a ubiquitous problem in science, and is particularly problematic in areas that depend heavily on computation and data analysis (see (Peng 2011)). For such work it is essential that installed software is identical to versions used in publication, in order to facilitate the reproduction of published data and the controlled manipulation or augmentation of these software systems. Unfortunately, this goal is often unattainable for a variety of related reasons: Research-oriented software may be hard to build and install due to unsatisfiable dependency constraints; non-trivial software may yield different results when built or used with different versions or variants of declared dependencies; on workstations and shared High Performance Computing (HPC) systems alike, it may be undesirable or even impossible to comply with version and variant requirements due to software deployment limitations. Moreover, It is unrealistic to expect users to manually recreate environments that match the system and binary substrate on which the software was developed. In the field of bioinformatics the above problem is exacerbated by the fact that data production technology moves extremely fast; existing software and data analysis workflows require frequent updates. Thus, it is paramount that multiple versions and variants of the same software can be automatically built, in order to ensure reproducibility of projects that are either in-progress, or are already published.

An important related issue is the reproducibility of workflows and pipelines across different machines. In addition to bioinformatics, many scientific fields require the researcher to prototype their code on local workstations with a custom software stack, and then later run it on shared HPC clusters for large data sets. The researcher must then be able to recreate their local environment on the cluster to ensure identical behavior. All of these concerns add to the burden on scientists, and valuable time that could be spent on research is wasted accommodating the limitations of system administration practices to ensure reproducibility. Even worse, reproducibility failures can be overlooked amid this complication, and publications could be accompanied with irreproducible analysis workflows or software. For these reasons, the scientific community in general -and fast evolving fields like bioinformatics in particular- need reliable and reproducible software package management systems.

In recent years, several tools have gained popularity among software developers and system administrators for wrapping Linux kernel features to accomplish process isolation, bind mounts, and user namespaces, or to deploy services in isolated environments (also called “containers”). Examples of such tools include: Docker, Singularity, and lxc. These tools are sometimes also proposed as solutions to the reproducibility problem (Peng 2011; Boettiger 2015), because they provide a way to ship an application alongside all of its runtime dependencies. This approach necessitates the use of file system images that are modified using imperative statements, e.g. to run a package manager inside a namespace, with the goal of embedding all dependencies in an opaque binary image. Containers and binary disk images alone do not make traditional tooling any more suitable for the purposes of reproducible science. Software deployment inside of the container is still subject to the well-known limitations of traditional package managers, such as intractable stateful behavior, time-dependent installation results, the inability to install and control more than a handful application or library variants of packages on the same system, to name a few. Container systems like Docker only shift the problem of reproducibility from the package level to the level of binary disk images, which is a much less useful level of abstraction. As such, they bring little more to the table than traditional virtual machine images, albeit with different trade-offs. We claim that reproducibility takes a more rigorous, declarative approach to software environment management and packaging itself. Other package and environment managers (such as Conda, EasyBuild, Spack) fail to take the complete dependency graph into account; instead, they make tacit assumptions about the deployment environment. As a result, it is much harder to understand and exactly reproduce an environment as neither the full complexity of the graph of transitive dependencies nor the configuration space is captured.

For all the above reasons, we propose functional package management -as implemented in GNU Guix- as a way to mitigate or obviate these problems by allowing us to *declare* the complete dependency graph of software packages (and all of their dependencies recursively). One important feature of this approach is that it allows for bit-by-bit reproducibility. To illustrate this, we created a set of analysis tools (or ‘pipelines’) for common genomics analysis data sets: RNA-seq, ChIP-seq, BS-seq and scRNA-seq (for the sequencing of RNA, Chromatin Immunoprecipitation, Bisulfite-treated DNA, and single-cell resolution RNA, respectively). Each pipeline has a complex and large graph of dependencies, and each graph is comprehensively declared as a GNU Guix package definition; the graph is then built reproducibly by relying on Guix package manager features. Note that these pipelines also represent production-level pipeline tools, rather than simply model examples -they come with a full set of features including alignment, quality check, quantification, assay specific analysis and HTML reports.

## Results

### Pipeline design and implementation philosophy

The pipelines provided here were designed with special focus on several key features: namely, that they be 1) easy to use, 2) easy to install, 3) easy to distribute, and, most importantly, 4) reproducible; all of which are inter-related constraints. Care was taken to ensure that all of the pipelines have a similar interface, so that familiarity with one pipeline would make for a gentler learning curve in learning to use the others. For the end-user, each pipeline has the same input types: a sample sheet and a settings file. The sample sheet contains information about samples such as names, labels, covariates etc. The settings file contains extra arguments related to the execution of the pipelines. The users can generally run pipelines as follows:

~~~
$ pigx [pipeline_name] [sample_sheet] -s [settings_file]
~~~

where [pipeline_name] can refer to any of the four pipelines: “rnaseq”, “chipseq”, “bsseq”, or “scrnaseq”. The resulting output provided to the users includes high quality reports and figures containing a standard set of results from basic analyses and data quality checks. Where appropriate, reports also contain certain interactive elements.

In implementing this toolset, one of our first design choices was to use a conventional build system, the GNU Autotools collection, to configure and build the pipelines as if they were first-class software packages in their own right rather than a mere collection of tools and “glue code”. Instead of assuming that a user will provide a suitable environment *at runtime*, the use of a build system allows us to capture the software environment *at configuration time*. This is achieved by explicitly checking for the presence of required tools in the build environment and recording their exact location in the pipeline’s configuration file. At runtime, the pipeline refers only to tools through the configuration file and does not assume the availability of dependent software in the global environment. Moreover, using a well-established build system makes it easy to package the pipelines for any package manager. We chose GNU Autotools over other build systems for two reasons: it does not require users to have a copy of the build system software as it compiles to shell code (which is highly portable), and it has been established long enough to implement a conventional and flexible build interface with well-known behavior even in somewhat unusual circumstances, such as the installation of files into unique prefixes as is done when building with GNU Guix.

Capturing the build-time environment alone is not enough to ensure reproducibility, nor is the use of a build system sufficient to make installation easy. Thus, our second design choice was to package the pipelines for the GNU Guix package manager. Like other package managers, GNU Guix allows users to install, upgrade and remove software without having to know the details of dependencies or the build procedure. Unlike traditional package managers, however, GNU Guix takes a rigorous, declarative approach to software environment management and packaging called functional package management. This approach takes into account the complete graph of dependencies and build-time configurations, and maximizes build reproducibility by building binaries in fully declared isolated environments. Packages are installed into paths with unique prefixes that are computed from the complete dependency graph, allowing for the simultaneous installation of different versions or variants of applications and libraries. With functional package management, a given software build will generally yield bit-identical files when the build is performed on different machines or on the same machine at different points in time, independent of the current state of the system (caveats to this generalization are discussed below).

We consider software reproducibility an important asset in controlled experimentation. Reproducing a software environment bit for bit is not a goal in itself, but it provides us with a foundation upon which we can perform precise changes to the environment and assess the impact of these changes. Without bit-for-bit reproducibility we cannot be certain of the nature and impact of differences in the software environment. While virtual machines or binary application bundles such as Docker images would be sufficient to freeze the state of our software environment, relying on these tools would forgo the ability to recreate that same environment from scratch; nor would it be possible to reason about the environment at the level of software packages. The approach of functional package management as implemented in GNU Guix preserves the relationships between software packages and ensures that differences to the environment can be accounted for.

A further design choice remained regarding the workflow management system, which would execute a series of tasks mostly in the form of scripts from different programming languages. For this purpose, we used SnakeMake (Köster and Rahmann 2012), which provides target-driven execution infrastructure similar to GNU Make but with Python syntax, along with useful features such as parallel execution on HPC scheduling systems. However, we would like to emphasize that the choice of workflow management system is not the most critical step for reproducibility, but rather the management of dependencies. The different pipeline stages are implemented with a workflow management system stitching together various bioinformatics tools; they are made configurable with the GNU Autotools and packaged with GNU Guix. This means they will be build-reproducible and can be installed via the one-liner:

~~~
guix package --install pigx.
~~~

## RNA-seq pipeline

### General Description of PiGx-RNA-seq Pipeline

PiGx RNA-seq provides an end-to-end preprocessing and analysis pipeline for RNA-seq experiments. The pipeline takes a set of raw fastq read files and the experimental design as described by the user, and produces differential expression reports with figures and tables of differentially expressed genes, as well as GO term analysis thereof. Furthermore, it provides quality control reports about the experiment. To use the pipeline, the user must provide two files: the sample sheet describing the samples and corresponding fastq files, and a settings file with configuration parameters related to the pipeline’s execution. The settings file lists, among other things, the location of a reference genome for alignment, a GTF file with genome annotations, and a transcriptome reference, as well as a list of desired differential expression analyses to be performed, specifying which samples to use as cases and controls --see package documentation here http://bioinformatics.mdc-berlin.de/pigx_docs/pigx-rna-seq.html for more details.

The pipeline can then be run with the command

~~~
$ pigx rnaseq [sample_sheet] -s [settings_file],
~~~

to generate the output -- which comes in several sequential steps (see figure 1).

**Figure 1.**
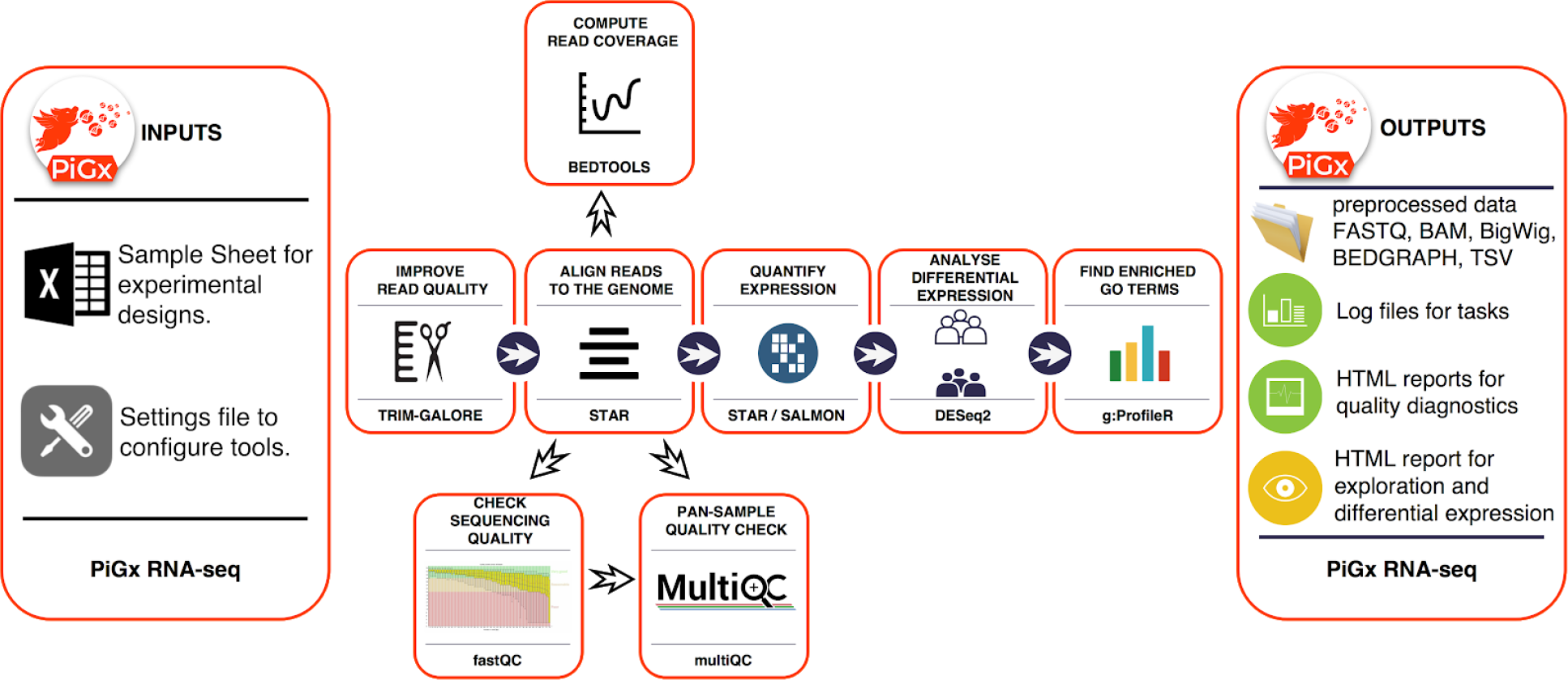
Workflow diagram of the PiGx-RNA-seq pipeline.

PiGx RNA-seq uses the reference genome and transcriptome provided by the user to produce indices using *STAR (Dobin et al. 2013)* and *Salmon (Patro et al. 2017)* respectively. It then uses *Trim Galore!* (Babraham 2018b) to trim low quality reads and remove adapter sequences before aligning the reads to the reference using *STAR*. At this point, PiGx RNA-seq uses *fastqc (Babraham 2018a)* and *MultiQC (Ewels et al. 2016)* to generate comprehensive quality control reports of the sequencing, trimming, and alignment steps. PiGx RNA-seq also uses *BEDTools (Quinlan and Hall 2010)* to compute the depth of coverage in the experiment and outputs convenient bedgraph files. Gene level expression quantification is obtained from *STAR*, and transcript level quantification using *Salmon*. The gene expression count matrix is then used to run differential expression analyses as specified by the user, using *DESeq2 (Love, Huber, and Anders 2014)* for statistical analysis and *g:ProfileR* (Reimand et al. 2007) for GO-term analysis. Each differential expression analysis produces a self-containing HTML report.

The differential expression reports produced are comprehensive, including sortable tables for differentially expressed genes for a detailed view, principal component analysis plots for a birds-eye view of the experiment, as well as MA and volcano plots. In addition, the reports include a section with GO term enrichment analysis.

### RNA-seq Use Case

The study by Hon *et al*. (2014) is motivated by several observations: DNA methyl-transferases (DNMTs) are the major mediators of cytosine methylation (producing 5-methyl-cytosine). 5hmC (5-hydroxy-methyl-cytosine) is a product of oxydation of 5mC’s, and the TET family of proteins mediate 5mC oxydation. It has been established that DNA demethylation consists of the sequence of chemical reactions that convert 5mC into 5hmC, which is subsequently converted into 5fC (5-formyl-cytosine) and 5caC (5-carboxyl-cytosine). Active enhancers are depleted for 5mC but are enriched for 5hmC marks (Rampal et al. 2014), suggesting that an interplay between DNMTs and TET proteins could determine the activity level of enhancers. Mutating DNMTs or TET proteins in mouse embryonic stem cells (mESCs) perturbs global DNA methylation status, however cells do not lose the ability to regenerate. Moreover, mutating TET proteins and perturbing the oxidation levels have previously been shown to skew the differentiation of mESCs. Based on these facts, the authors address the following question: Can the skewed differentiation in mESCs be explained by deregulated balance of 5mC / 5hmC levels at active enhancers following the loss of activity of TET proteins?

The authors of the above study use TAB-Seq, Bisulfite-Seq, ChIP-seq and RNA-seq methods to profile genome-wide methylation, demethylation, histone modifications and gene expression levels to address these questions. They find that *Tet2* has the biggest role in enhancer demethylation in mESCs. Deletion of *Tet2* leads to enhancer hypermethylation, which in turn reduces enhancer activity. The reduced enhancer activity leads to a disruption in the activation of more than 300 genes in the early stages of differentiation, however the activity levels of these genes are restored to wild-type levels at the later stages of differentiation. Reduced enhancer activity followed by delayed gene activation explains the skew observed in mESC differentiation.

The authors of the above study profile the transcriptomes of mESCs as they differentiate into neural progenitor cells (NPCs) within a six day period. They quantify gene expression levels of wild-type, *Tet1* -/- and *Tet2* -/- cells on day zero, day three, and day six and sequenced two biological replicates per sample. Thus, they obtained 18 samples in total (3 genotypes x 2 replicates x 3 days). In figure 5 of the original manuscript, the authors summarise the results of the RNA-seq analysis. Here, we use the PiGx-RNA-seq pipeline to pre-process the raw fastq files downloaded from the GEO archive (GEO accession: GSE48519), map the reads to the *Mus musculus* genome (GRCM38 (mm10) build), and finally quantify the expression levels of genes using both Salmon (Patro et al. 2017) and STAR (Dobin et al. 2013). We then use DESeq2 (Love, Huber, and Anders 2014) to perform multiple differential expression analyses as described in the original publication. Based on the processed and normalized count tables and differential expression analysis results produced by the PiGx pipeline, we have written a small custom script to reproduce the panels in figure 5 of Hon *et al*. In order to reproduce this figure, we needed to perform seven differential expression analyses as described in Table 1. HTML reports for each differential expression analysis (based on read counts computing using STAR) can be found here: http://bioinformatics.mdc-berlin.de/pigx/supplementary-materials.html.

**Table 1.** Differential expression analyses performed by PiGx-RNA-seq.

| Analysis | Case Sample | Control Sample | Description |
| --- | --- | --- | --- |
| tet2_diff_day3 | day3_tet2_KO | day0_tet2_KO | *Tet2* *-/-* cells on day 3 are compared to *Tet2* *-/-* cells on day 0. |
| tet2_diff_day6 | day6_tet2_KO | day0_tet2_KO | *Tet2* *-/-* cells on day 6 are compared to *Tet2* *-/-* cells on day 0. |
| WT_diff_day3 | day3_WT | day0_WT | Wild-type cells on day 3 are compared to wild-type cells on day 0. |
| WT_diff_day6 | day6_WT | day0_WT | Wild-type cells on day 6 are compared to wild-type cells on day 0. |
| tet2_vs_WT_day0 | day0_tet2_KO | day0_WT | *Tet2* <sup>-/-</sup> cells on day 0 are compared to wild-type cells on day 0. |
| tet2_vs_WT_day3 | day3_tet2_KO | day3_WT | *Tet2* <sup>-/-</sup> cells on day 3 are compared to wild-type cells on day 3. |
| tet2_vs_WT_day6 | day6_tet2_KO | day6_WT | *Tet2* <sup>-/-</sup> cells on day 6 are compared to wild-type cells on day 6. |

Having performed the above analysis, we first took a global look at how all sequenced samples cluster. Using a table of TPM (transcripts per million reads) counts generated by Salmon at the gene level, we selected the top 100 most variable genes and plotted a heatmap of all the samples using pheatmap package (Kolde 2018). We observed that the samples mainly cluster by the differentiation stage rather than genotype, which confirms the authors’ findings (figure 2A). Next, again using the same TPM counts table, we plotted the expression levels of a select list of genes (*Nes6, Pax6, Sox1, Tet1, Tet2, Tet3, Slit3, Lmo4, Irx3*) on day 0, day 3, and day 6 (figure 2B). The changes in the expression levels of these genes perfectly match the patterns as described by Hon et al. At this point the authors recognise that some neural marker genes such as *slit3* and *lmo4* show discordant expression patterns between WT and *Tet2* -/- samples particularly on day 3, which are restored back to WT levels on day 6. The authors then investigated whether such a delayed induction mechanism can be observed globally. It was shown that the percentage of genes that are differentially expressed in both *Tet2* -/- and WT cells (compared to the undifferentiated samples of the corresponding genotypes on day 0), is significantly higher on day 6 than on day 3. We also observe a similar pattern, however the difference we observe is somewhat reduced. Our findings are reproduced based on gene counts quantified by both STAR and Salmon (figure 2C).

**Figure 2.**
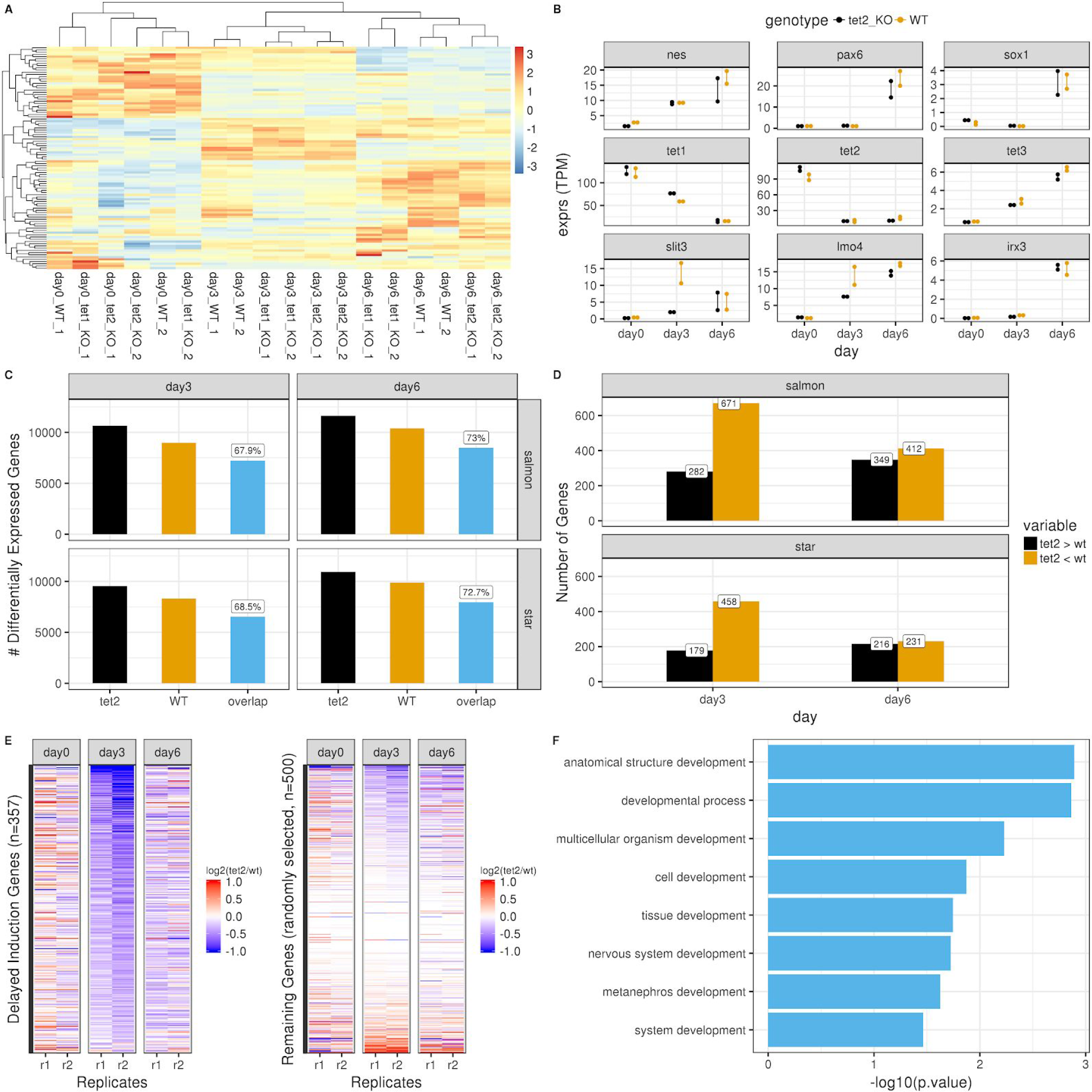
Reproduction of figure 5 from (Hon et al. 2014) using datasets processed by PiGx-RNA-seq pipeline. **A)** Hierarchically clustered heatmap of the top 100 most variable genes across all samples (transcripts per million (TPM) aggregated on the gene level, produced with Salmon). Each row represents a gene and each column represents a sequenced sample (See Table 1 for descriptions of the samples). The expression values are scaled by ‘row’. **B)** Changes in the expression levels of a selected list of genes throughout differentiation period on day 0, day 3, and day 6. The y-axis shows the normalised expression levels (TPM at gene-level). The expression patterns of samples with *Tet2* -/- background are depicted in black and wild type background in orange. **C)** Abundance of differentially expressed genes (adjusted p-value < 0.1) (on y-axis) when comparing samples on day 3 or day 6 with the samples on day 0 with corresponding genotypes (*Tet2* -/- or wildtype). The bar labeled ‘overlap’ represents the number of differentially expressed genes in both genotypes. The percentage is calculated by dividing the value of ‘overlap’ with the value of *Tet2*. The results are reproduced by both Salmon-based gene-level read counts (top row) and STAR-based gene-level read counts (bottom row). **D)** Genes that are up-regulated (induced) in wild-type samples on day 3 (or day 6) compared to wild-type samples on day 0, are intersected with genes that are differentially expressed between wild-type samples and *Tet2* -/- samples at the same stage of differentiation, and classified as *‘Tet2* > wt’ (the gene is up-regulated in the *Tet2* -/- sample moreso than in the wild-type sample) or *‘Tet2* < wt’ (the gene is upregulated in *Tet2* -/- sample less than in the wild-type sample). The plot is reproduced using both Salmon-based gene counts and STAR-based gene counts. **E)** Heatmaps for delayed induction genes (on the left) and 500 genes randomly selected from the remainder (on the right). The colors of the heatmap represent the log_2_ scale ratio of normalised expression value (gene-level TPM counts obtained using Salmon) of each delayed induction gene between *Tet2* -/- sample and the wild-type sample of the corresponding replicates (r1: replicate-1, r2: replicate-2) on the corresponding stages of differentiation (day 0, day 3, and day 6). The rows of the heatmap are ordered in increasing order based on the average values of the two replicates on day 3. The color scales range between -1 and 1 before reaching saturation. **F)** Top GO terms for biological processes (on the y-axis) enriched among the delayed induction genes. The GO terms are detected using g:ProfileR tool (Reimand 2016). The resulting terms are filtered for p-value<0.05 and further filtered for the keyword ‘development’. On the x-axis, the p-values are depicted at log_10_ scale.

In figure 5F of the original publication, the authors take a closer look into the list of discordantly induced genes on day 3 in *Tet2* -/- samples. There it is shown that the majority of the genes that get induced in WT samples by day 3, don’t get induced in the *Tet2* -/- samples as highly as they do in the WT samples. On the other hand, these numbers are comparable on day 6. We also observe the same difference and reproduce the findings using both Salmon and STAR-based gene counts (figure 2D). This suggests that there must be a list of genes that get activated in WT, but lag behind in *Tet2* -/- samples at the early stage of differentiation, however they catch up later with the WT levels. The authors call these genes ‘*delayed induction genes*’ and find 333 genes that fit such a description. In figure 5G, the authors show the relative expression of these genes in *Tet2* -/- samples compared to WT samples throughout differentiation and compare it to the remaining list of genes in the genome. We have successfully reproduced the same patterns based on 357 delayed induction genes detected by Salmon-based gene counts (282 genesdetected by STAR-based gene counts) (Figure 2E). In figure 5H, the authors show the most significant GO terms enriched for the delayed induction genes. Although we don’t observe the same set of terms as reported by the authors, we found seven development-related GO terms including ‘tissue development’ and ‘nervous system development’ as enriched terms (figure 2F).

## ChIP-seq pipeline

### General Description of PiGx-ChIP-seq Pipeline

PiGx ChIP-seq is an end-to-end processing and analysis pipeline for ChIP-seq experiments. From the input fastq files, the pipeline produces sequencing quality control, ChIP quality control, peak calling, and IDR (Q. Li *et al*. 2011) estimation. PiGx ChIP-seq also prepares the data for visualization in a genome browser. The pipeline execution is highly customizable - the user can specify which parts of the pipeline to execute, and which parameter settings to use. As in the other pipelines, to use PiGx ChIP-seq, the user must provide two files: a sample sheet containing the names of the fastq files with a descriptive label, and a settings file. The settings file contains the locations of the reference genome, and the GTF file with genome annotations, as well as a list of configurations for each executable step. Upon completion, the user is provided with quality reports, and all of the pre-processed data, which substantially facilitates downstream analysis and visualization.

PiGx ChIP-seq pipeline aligns the reads to the genome using Bowtie2 (Langmead and Salzberg 2012), does peak calling using MACS2 (Zhang et al. 2008), calculates the irreproducibility rate and outputs a series of quality statistics, such as: GC content, strand cross correlation, distribution of reads and peaks over annotated genomic features, and clustering of samples based on their similarity (Landt et al. 2012). The pipeline also produces UCSC Track hub for exploration of the dataset. The purpose of the pipeline is to improve the routine processing steps for ChIP-seq experiments and enable the user to focus on data quality control and biologically relevant data exploration. The pipeline heavily depends on Bioconductor (Huber *et al*. 2015) packages such as GenomicRanges (Lawrence et al. 2013) and Genomation (Akalin *et al*. 2015) for annotating peaks and summarizing ChIP-seq scores over regions of interest.

**Figure 3.**
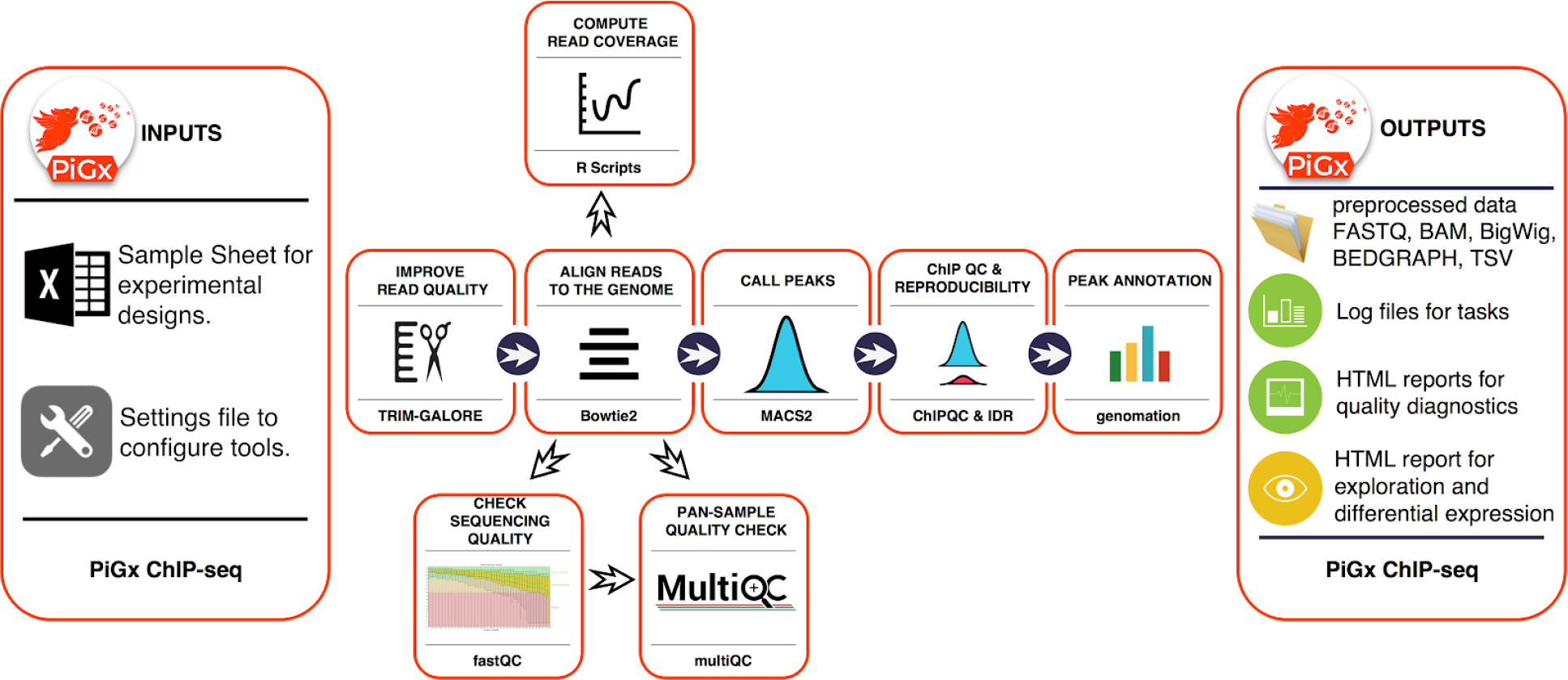
Workflow diagram for ChIP-seq pipeline

### ChIP-seq Use Case

For consistency, we applied the ChIP-seq pipeline to data from the same study as in the section “RNA-seq Use Case” above (Hon *et al*. 2014); for the biological underpinnings of this experiment, please see the description provided there. Figure 4 shows part of the ChIP-seq quality control output performed on untreated, wild type ChIP samples, of various activating and repressing histone marks, and the corresponding input samples. One standard procedure is to validate the consistency of results with known biological priors, in order to quickly find samples with outlying properties, and to discover batch effects. For example, figure 4A shows the expected clustering of repressive (H3k27me3, H3k9me3) and activating (H3k4me3, H3k4me1, H3k27ac, and H4k36ac) histone marks. Upon closer inspection, however, it becomes clear that the activating histone marks cluster by their corresponding *batches*, and not by their biological functionality. Figure 4B shows the cross-correlation between the signal on the plus and minus genomic strands, shifted within a defined range (usually 1 - 400 nucleotides). The maximum intensity in each row indicates the average DNA fragment size in each corresponding ChIP experiment. Large discrepancies in the cross correlation profile, between experiments, can indicate problems with fragmentation, fixation, or chromatin immunoprecipitation. The figure shows that most of the samples have an average fragment size between 100 - 150 bp. One of the H3k27me3 replicates, however, shows aberrant fragment size profile (second sample in the plot). Upon visual inspection, the sample had extremely low signal to noise ratio and the peak calling resulted in zero enriched regions. Such samples should either be repeated or omitted from the downstream analysis. Figure 4C represents the relationship between the GC content of one kilobase genomic bins and the ChIP signal; this plot is used as a diagnostics tool for enrichment of fragments with extreme nucleotide content (enrichment of fragments with GC content strongly deviating from the genomic mean), which can indicate problems with PCR-based fragment amplification, and chromatin immunoprecipitation. Figure 4D represents the distribution of reads over functional genomic features. It is used to observe whether the experimental results conform to known expectations, based on previous experiments - i.e. H3k4me3 should show strong enrichment over transcription start sites, while the H3k36me3 should show an enrichment over exonic and intronic regions. Non-conforming experiments can indicate a weak ChIP, or antibody cross reactivity with unexpected epitopes. Figure 4 represents just a subset of quality control metrics implemented as a standard output from the PiGx- ChIP-seq pipeline. The full set can be found here: http://bioinformatics.mdc-berlin.de/pigx/supplementary-materials.html

**Figure 4.**
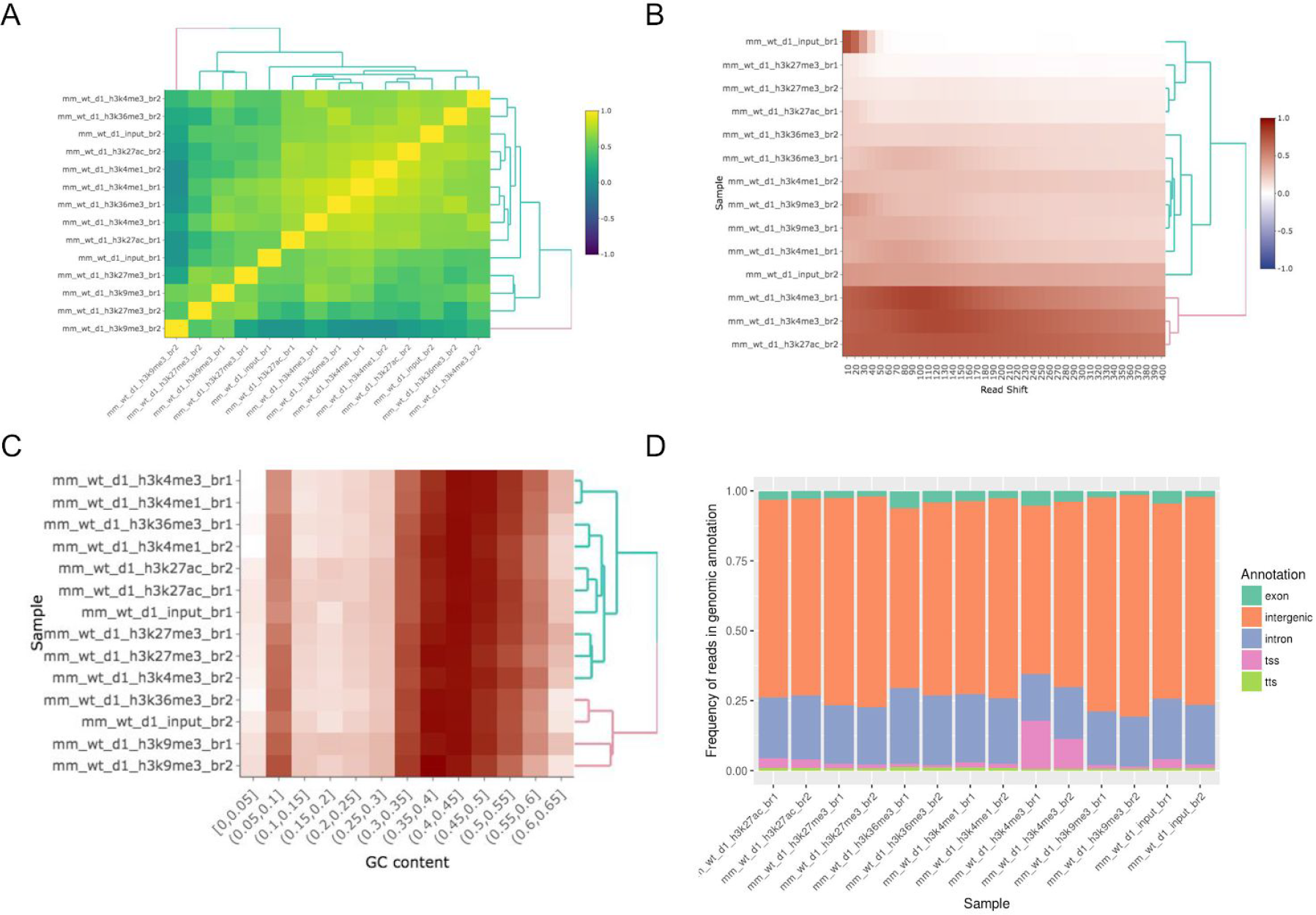
Example ChIP-seq quality control output. **A)** Clustering of samples based on correlation of normalized ChIP reads in one kilobase bins. **B)** Cross correlation between coverage profiles on Watson and Crick strands, shifted by the amount specified on the x axis. **C)** Relationship between read count and GC content in 1 kb bins. **D)** Distribution of reads in functional genomic features.

## BS-seq pipeline

### General description of the PiGx BS-seq pipeline

PiGx BS-seq is a bisulfite sequencing processing pipeline used to detect genome-wide methylation patterns and to perform differential methylation calling for case-control settings. It produces individual reports for each sample provided by the user, in addition to differential-methylation reports for arbitrarily many pairs of treatment conditions provided by the user. PiGx BS-seq uses *Trim Galore! (Babraham 2018b)* to trim reads for adapter sequences and quality, and *fastqc (Babraham 2018a)* for quality control (both before and after trimming). PiGx BS-seq produces *GA*- and *CT*- converted versions of the reference genome, if necessary, using bismark_genome_preparation (Krueger and Andrews 2011). Reads are then mapped to the reference using Bowtie2 (Langmead and Salzberg 2012), before being sorted by location in the genome and filtered for uniqueness using samtools (Krueger and Andrews 2011; H. Li *et al*. 2009). The corresponding reports and .bam files for each of these steps are saved to their respective directories.

As in the other pipelines, to use PiGx BS-seq, the user must provide two input files: a sample sheet containing the paths to the fastq files with a descriptive label, and a settings file. The pipeline is robust to paired-end or single-end input data, and processing of each case is initiated automatically, based on whether the user supplies only a single input file, or a pair of files, for each sample. The settings file contains the locations of the reference genome, among other directories, as well as a list of configuration steps for each executable step. The pipeline can then be run with the command:

~~~
$ pigx bsseq [sample_sheet] -s [settings_file],
~~~

Post-mapping analysis steps performed automatically by PiGx BS-seq include tabulation of the fractional methylation of CpG sites, the segmentation of genomic methylation patterns across the genome, and the selection of differentially methylated sites between pairs of treatments provided in the settings file above. Furthermore, the final reports include genomic annotation of differentially methylated regions and methylome segments. A single execution of the pipeline can perform differential methylation analysis between a sample and arbitrarily many references; each comparison will have its own dedicated report, in addition to the final report for the sample itself. For traceability, direct links to input files, and various execution tools are saved directly within the output folder. Finally, a copy of the full methylome for each sample is also saved in BigWig (.bw) format, compatible with visualization in an online genome browser.

**Figure 5.**
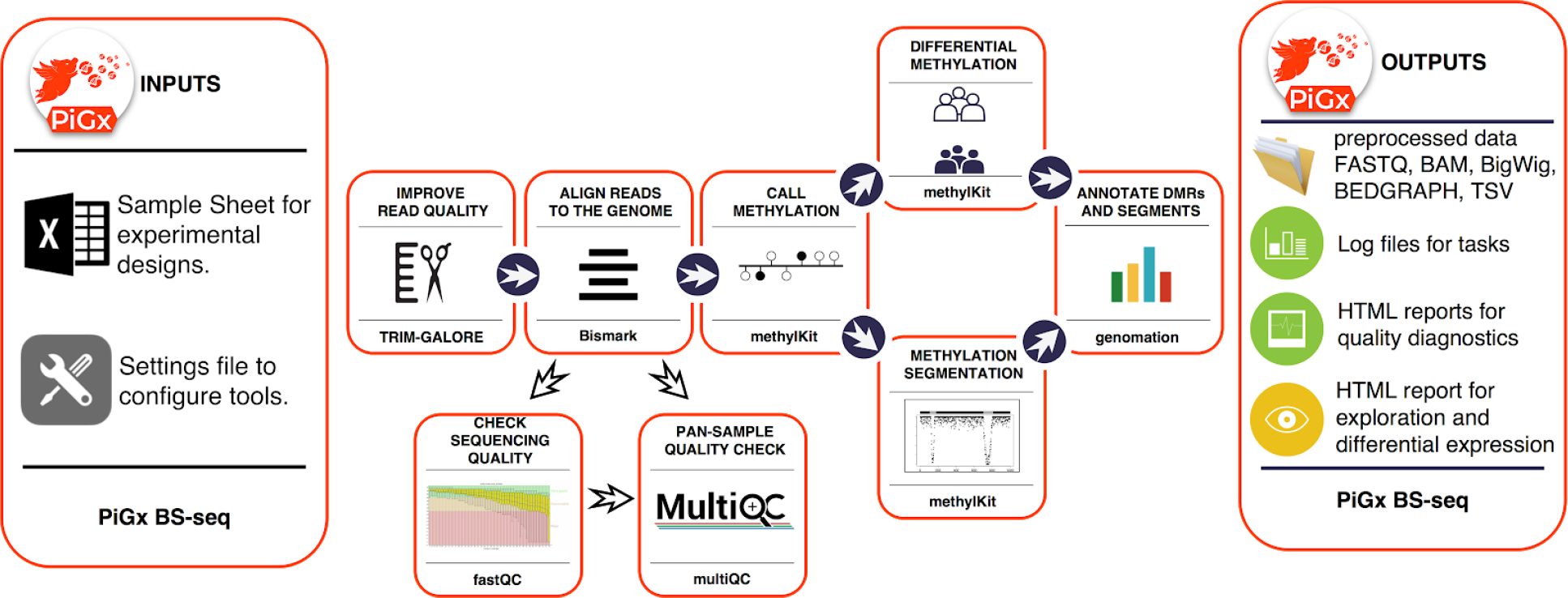
Workflow diagram for PiGX BS-seq pipeline

### BS-seq Use Case

We applied the BS-seq pipeline to data from embryonic stem cells in mice, comparing wild type and *Tet2* deletion experiments (accessions SRX317877, and SRX317883 respectively). These data sets derive from the same study as was used for controlled comparison in the section “RNA-seq Use Case” above (Hon *et al*. 2014); for a biological description of this experiment, please refer to that section. HTML reports for each of the performed analyses can be found here: http://bioinformatics.mdc-berlin.de/pigx/supplementary-materials.html

Figure 6 shows a standard set of data analysis metrics generated automatically by the pipeline. For example, methylation levels near the promoter region of a list of annotated genes for each sample are shown in figures (A) and (B). For generality, figure 6 averages over all known genes; the user may freely probe for more specific results by supplying any arbitrary set of genes under investigation (in the absence of such an annotation file, this figure is simply omitted from the final report). A coarse map of the genome is provided in (C), which, for some datasets, may serve to highlight differential methylation localized to particular regions or chromosomes. In this particular use-case it is more useful as a null control showing that these regions are uniformly distributed throughout the genome. In addition, a histogram for differential methylation status of CpGs throughout the genome is provided in (D) using the same colour-code as in (C). The methylation differences of hyper-methylated, hypo-methylated and non-differentially methylated CpGs are shown as histogram with the color-code as in Figure 6C. The latter is shown as a distribution of methylation differences deemed to be not statistically significant (in black); since these are generally far more numerous than the former, the two curves are normalized independently. Note also that since these curves represent *relative* distributions, the vertical axis is of arbitrary units and tick marks are omitted. Finally, a screenshot of data-visualization from the genome browser (Robinson et al. 2011; Thorvaldsdóttir, Robinson, and Mesirov 2013) is provided in (E), here, regions of interest can be inspected manually at arbitrary precision.

**Figure 6.**
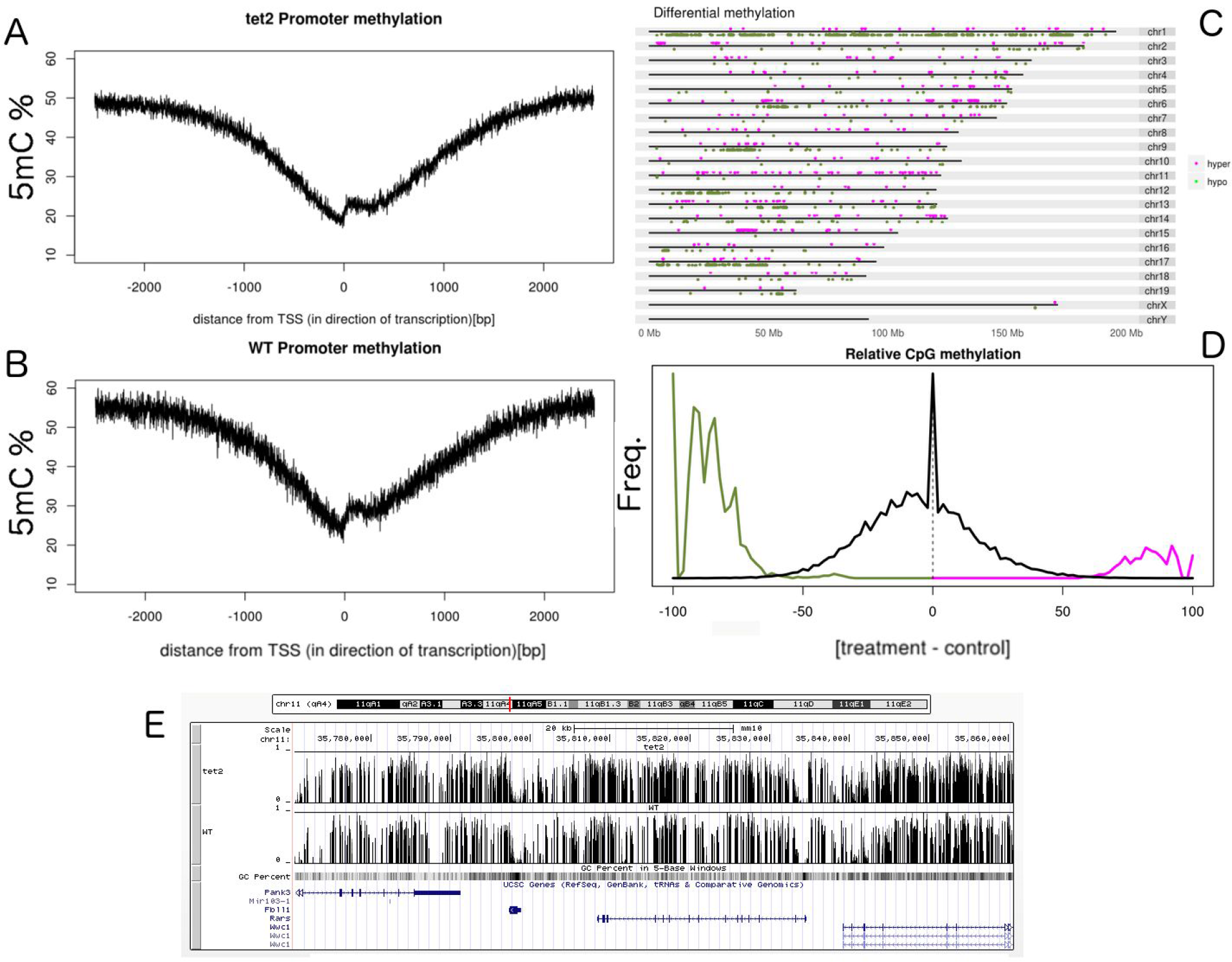
Output from the PiGx BS-seq pipeline. **(A,B)**: average CpG methylation throughout the promoter regions of the mm10 genome for *Tet2* -/- and WT respectively. **C)** Whole-genome map of differentially methylated CpGs, with colour-code to indicate hyper- and hypo- methylation of the treatment (*Tet2* -/-) relative to the control (Wild-type). **D)** Histogram of the difference in average CpG methylation between *Tet2* -/- and wild-type. For differentially-methylated cytosines, colors are consistent with (C), while CpGs with statistically insignificant difference in methylation are provided in black. Normalization of these two curves is performed independently (since the latter are generally far more numerous than the former), and the graph conveys only relative proportions (thus, as the absolute y-axis is of arbitrary scale, units and tick marks are omitted). **E)** Screenshot of the genome browser using bigwig data from PiGx; here the data can be examined in much finer detail than in C).

## scRNA-seq pipeline

### General description of the PiGx scRNA-seq pipeline

Single cell RNA-seq is an extremely powerful technology, that is becoming increasingly prevalent in biological studies. The rapid development of UMI based methods, along with droplet based cell separation (Macosko et al. 2015; Klein et al. 2015), has enabled even simple experiments to quantify expression in several tens of thousand of cells. **PiGx scRNA-seq** is a pipeline for pre-processing of UMI based single-cell experiments. The purpose of the pipeline is to enable seamless integration and quality control of multiple single cell data sets. The pipeline works with minimal user input. As in the other pipelines, the user must provide a sample sheet with a basic experimental description, and a settings file which defines, among other parameters, the location of the input data and reference sequence and annotation. The pipeline can then be run with the command:

~~~
$ pigx scrnaseq [sample_sheet] -s [settings_file]
~~~

The pipeline does preliminary read processing, maps the reads with the STAR (Dobin et al. 2013) aligner, and assigns reads to gene models. It also separates cells from background barcodes (Alles et al. 2017), and constructs digital expression matrices for each sample (each saved in loom format); loom files from all samples are then merged into one large loom file using the loompy package (Linnarsson 2018). The expression data are subsequently processed into a SingleCellExperiment (Aaron Lun and Risso 2018) object. SingleCellExperiment is a Bioconductor class for storing expression values, along with the cell, and gene data, and experimental meta data in a single container. It is constructed on top of hdf5 file based arrays (Pagès 2018), which enables exploration even on systems with limited RAM (random access memory).

During the object construction, the pipeline performs expression normalization, dimensionality reduction, and identification of significantly variable genes. Then, it classifies cells by cell cycle phase and calculates the quality statistics. The SingleCellExperiment object contains all of the necessary data needed for further exploration. The object connects the PiGx pipeline with the Bioconductor single cell computing environment, and enables integration with state of the art statistical, and machine learning methods (scran (A. T. L. Lun, McCarthy, and Marioni 2016), zinbwave (Risso et al. 2018), netSmooth (Ronen and Akalin 2018), iSEE (Aaran Lun et al. 2018), etc.).

The pipeline produces an HTML report containing quality controls, labeled by input covariates, which can be used for detecting batch effects.

### scRNA-seq Use Case

To showcase the capabilities of PiGx scRNA-seq, we ran the pipeline on isolated single nuclei from the mouse brain (Hu *et al*. 2017). In this study, the authors developed a gradient-based method for nucleus separation, and used it in combination with Drop-seq to profile the transcriptomes of more than 18,000 single nuclei. Figure 8 shows a part of the quality control output from the PiGx scRNA-seq pipeline. Figure 8A shows the per sample number of total and uniquely mapped reads. Figure 8B visualizes the cells on the first two principal components. The color gradient corresponds to the number of detected genes per cells. The figure shows that the total number of detected genes strongly correlates with the first two principal components. Figure 8C is analogous to figure 7B of the original publication, with the color scheme representing labeling each cell with its respective stage of the cell-cycle. Thus, figure 8C shows that the first two principal components correlate with the stage of the cell cycle. The heatmap in figure 8D shows scaled normalized expression values for genes that contribute the most to the first principle component. High read-count variability in a small number of genes drives the variation around the first principle component. The column-wise annotations show that the variation is driven mainly by cells in the G1 phase of the cell-cycle from the second biological replicate. The HTML report for this analysis can be accessed here: http://bioinformatics.mdc-berlin.de/pigx/supplementary-materials.html

**Figure 7.**
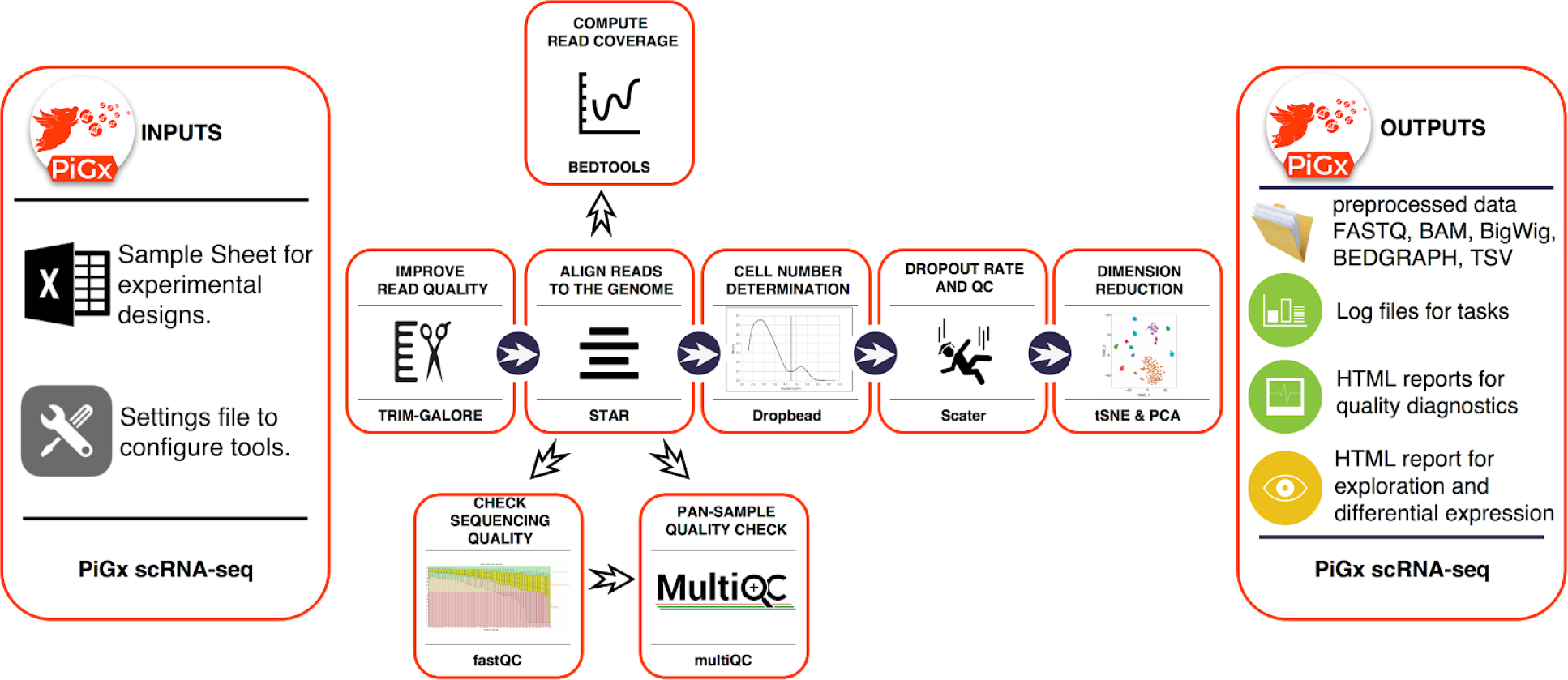
Workflow diagram for PiGx-scRNA-seq pipeline.

**Figure 8.**
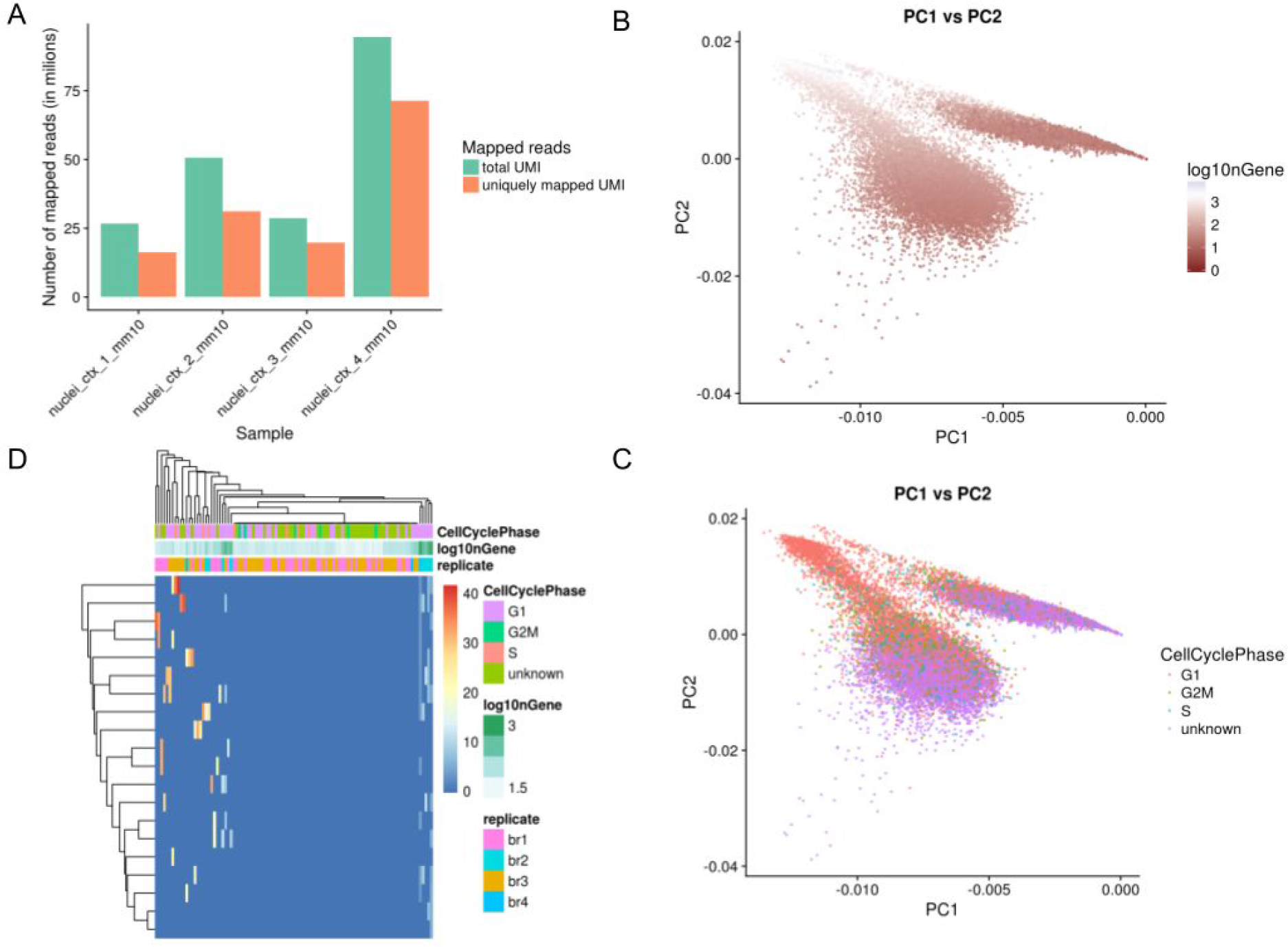
Sample output from the PiGx scRNA-seq pipeline. **A)** Abundance of total uniquely mapping UMIs per sample. **B)** Visualization of cells on the first and second principal component calculated from the normalized expression values. The gradient shows the total number of UMIs per cell. **C)** Same data representation as in B, but colored based on the cell cycle assignment. Cell cycle was assigned using the cyclone function from the scran Bioconductor package (A. T. L. Lun, McCarthy, and Marioni 2016). **D)** Expression heatmap of genes contributing most to the first principle component. Genes are ordered in rows, while cell are in columns. Color bars above the heatmap show relevant experimental variables.

## Reproducibility metrics of the pipelines in different systems

We define the complete software environment needed for each of the pipelines using Guix package definitions. These package specifications not only outline the immediate dependencies of the pipelines, but extend to the full software stack recursively. The dependency graph is rooted in a handful of bootstrap binaries. Apart from these binary roots, every application or library in the graph is built from source. Guix ensures that packages are built in an isolated environment in which nothing but the specified dependencies are available. This is a precondition for bit-reproducible builds, i.e. repeatable package builds that yield the very same binary output for the same set of inputs. Under ideal circumstances, a Guix specification for the complete dependency graph and the set of all source code would be sufficient to exactly reproduce the very same binaries of the pipelines presented in this paper.

Unfortunately, there are additional obstacles to bit-reproducibility that cannot be avoided purely by the functional package management model. Examples for sources of irreproducibility in build artefacts include embedded timestamps, non-deterministic sorting of strings, non-deterministic compiler output, and the like. While some of these obstacles can be removed by deliberate patching of compilers or applications, others are harder to diagnose and can thus lead to failure to reproduce the same arrangement of bits in independent builds, be that on the same machine at different points in time or on different systems.

To estimate the level of bit reproducibility in our pipelines, we checked out version v0.14.0-3597-g17967d1 of GNU Guix, repeatedly built the pipeline packages and their direct dependencies on three different systems (an office workstation, a virtual machine, and a build farm consisting of 20 heterogeneous build nodes), and recorded the hashes of the produced package trees. Whenever the hashes of any two builds differed we looked at the exact differences with diffoscope (https://diffoscope.org/). Upon closer inspection we identified a number of common issues in non-deterministic builds, such as timestamps embedded in compiled binaries and text files, or randomized file names in files generated by test suites.

Python dependencies are of particular note here, because they are generally not reproducible due to the fact that the byte compiler records the timestamp of the source file in the compiled binary. This means that all compiled Python files will differ when they are compiled at different points in time. (This problem will be addressed in the upcoming Python 3.7, which will implement PEP 552 for deterministic compilation.) To avoid this problem and increase the number of packages that could be made reproducible, we patched our variant of Python 3.6 such that it resets the embedded timestamp in compiled files to the Unix epoch. This allowed us to greatly increase the number of fully bit-reproducible packages. As can be seen in Table 2, only a total of 8 out of 355 packages (or only about 2.2%) were not bit-reproducible for as yet unknown reasons.

**Table 2.** Number of dependent packages and their reproducibility status. See Table 3 for more details about packages with minor problems.

| Package | Not reproducible | Minor problems | Reproducible |
| --- | --- | --- | --- |
| pigx-bsseq | 2 | 2 | 167 |
| pigx-chipseq | 7 | 9 | 236 |
| pigx-rnaseq | 7 | 9 | 211 |
| pigx-scrnaseq | 6 | 8 | 218 |
| All pipelines | 8 | 9 | 338 |

**Table 3.** Table of packages with minor reproducibility problems and the magnitude of irreproducible files.

| Package | Magnitude | Notes |
| --- | --- | --- |
| r-minimal | 2 bytes | non-deterministic line break |
| python | ~ 6% | timestamp byte in header of bytecode files |
| python-matplotlib | ~ 1.7% | single file difference |
| python-pycparser | ~ 3% | single file with timestamp |
| python-cffi | ~ 1.8% | recorded random test file names |
| python-numpy | < 0.5% | six bytecode files differ |
| python-simplejson | 2 bytes | two files have single byte differences |
| gtk+ | < 1% | single file (icon cache) |
| glib | < 0.1% | single file difference |

Figure 9 visualizes the degree of bit-reproducibility for the direct dependencies of each of the individual pipeline packages. Dependent packages whose files differed compared to builds on other systems fell either in the category of “minor problems” or “not reproducible”, dependent on the source and magnitude of non-determinism. The exact dependency counts for each category and pipeline package are listed in Table 2. A comprehensive list of all dependent packages that were categorized as having “minor problems” is contained in Table 3. This table shows that the reproducibility problems of these packages are of negligible magnitude and could be corrected with minor patches to the package definitions in Guix.

**Figure 9.**
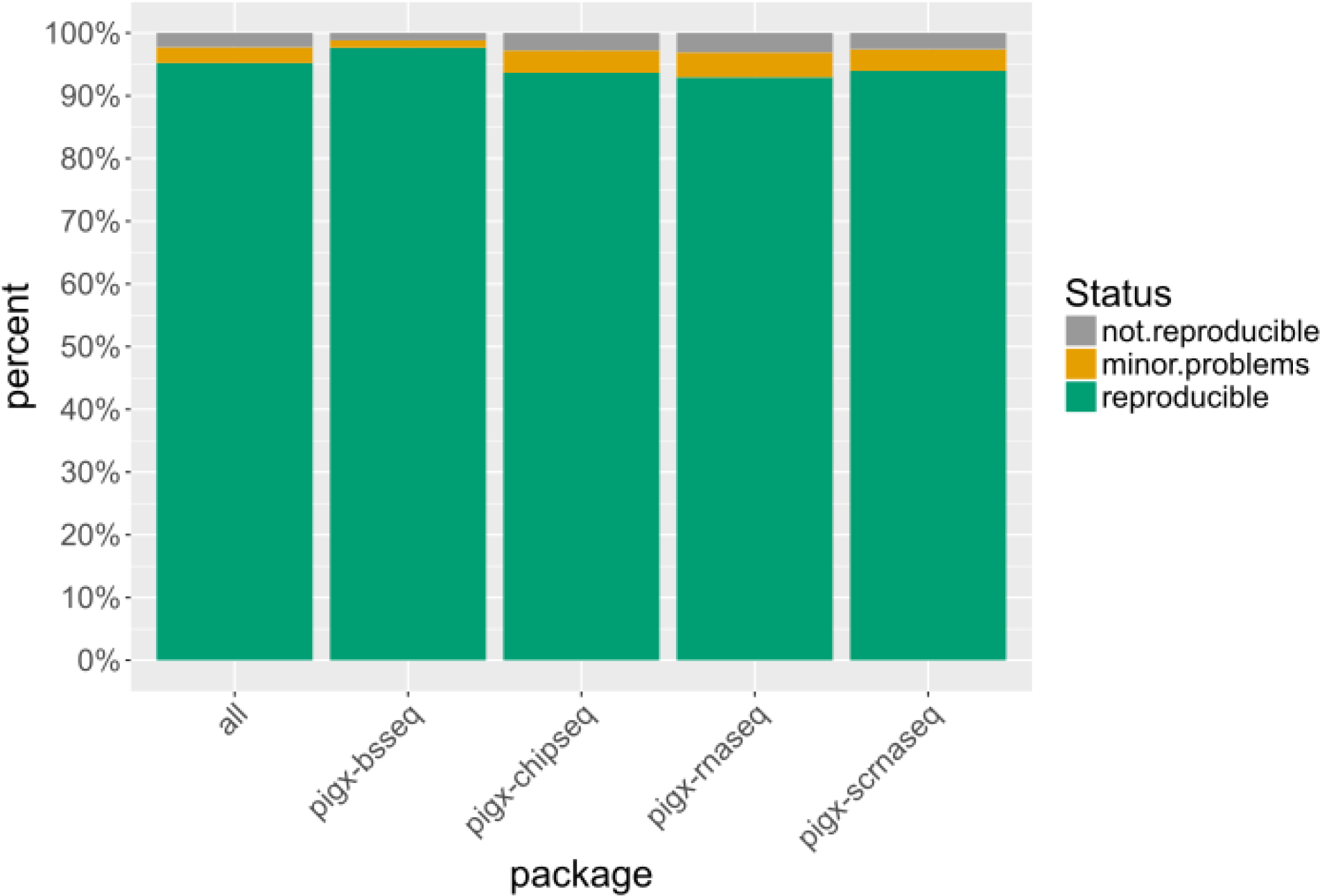
Percentage of directly-dependent packages building in a bit-reproducible fashion across different systems for each of the pipelines.

## Alternative ways to install the pipelines

We provide a generated application bundle containing all pipelines for use with Docker. The Docker image was generated by exporting the "closure" (i.e. the package and all packages it references, recursively) of the pigx package from the declarative Guix package definition instead of iteratively modifying a base image containing a GNU+Linux operating system in a series of imperative steps. The Docker image is merely a translation of a functional description of the desired environment; consequently, it is independent to global state, such as the contents of third-party package repositories or build time. The Docker image can be obtained at https://hub.docker.com/r/bimsbbioinfo/pigx/

Since the pipelines use the well-known GNU build system as implemented by the Autotools suite, the pipelines can be configured and built in any environment providing the required dependencies. The portable configure script detects and records references to needed software in the environment and reuses them at runtime using their absolute file names. Any package manager (such as Conda) can be used to fashion such a build-time environment. With regards to reproducibility, however, we recommend that a package manager be used that can provide separate, immutable, and uniquely prefixed environments to ensure that references to tools that are recorded at configuration time are identical to the variants that are used at runtime.

## Discussion

Computation is becoming an increasingly essential part of the biological sciences as the field becomes more data intensive. The diversity and amount of data requires a multitude of tools being used for analysis. Thus, the published software or workflows often come with complex dependencies. Even if sensible guidelines (e.g. “Software with Impact” 2014), such as sharing code online and providing documentation, are employed, sometimes it is impossible to recreate the software used for analysis. Providing the code and documentation alone does not guarantee reproducibility or usability, nor do Docker containers completely remedy this problem. We propose GNU Guix and principled pipeline-as-software implementation as a solution to reproducibility problems in complex bioinformatics workflows. Here, we demonstrated the utility and the reproducibility of PiGx pipelines for genomics data analysis using GNU Guix.

Our decision to treat pipelines as first-class software packages and to adopt a conventional build system with Autotools made it possible to reduce the installation of complex software environments to a simple one-line command. By recording the exact locations of runtime dependencies of the pipeline packages during the configuration stage, we were able to eliminate ambiguity at runtime. When configuring the pipeline packages in an environment that ensures that different versions or variants of applications and libraries are stored in unique locations (such as an environment provided by GNU Guix), recording the exact location of dependencies at *configuration time* allows us to reproduce the detected environment at *runtime*.

We have shown that with a recursive definition of software dependencies using the framework provided by the functional package management paradigm as implemented in GNU Guix, it is possible to fully and exhaustively describe complex real-world bioinformatics software environments. The software environments were fully specified at the level of declarative, stateless package abstractions instead of using an imperative, stateful approach. We have also shown that the principled declarative approach to the management of software environments lays a solid foundation for bit-reproducibility. The higher-level definitions of software environments can be translated in an automated fashion to lower-level application bundles such as Docker images. In contrast with container systems like Docker or Singularity, Guix encloses the complete software environment and enables users to transparently rebuild it reproducibly from source without having to trust a binary application bundle. Due to referential transparency, binaries in Guix can only be the result of their corresponding sources.

Functional package management as implemented by GNU Guix significantly reduces the complexity of, and lowers the barrier to, managing bit-reproducible software environments. Users are freed from menial bookkeeping tasks such as keeping track of the origin of package binaries, the time of installation, the order of installation instructions, the state of the operating system at the time of installation, or any other runtime state. As far as users are concerned, it is enough to know the names of the packages that should be installed (in our case, simply “pigx”) and the current version of Guix; everything else such as source code provenance tracking, dependency management, package configuration, and compilation in isolated environments is handled by Guix. The guarantees provided by Guix enable users to contemplate obstacles to experimental reproducibility beyond the software environment, such as sources of non-determinism at *runtime*.

In our attempts to analyze the degree of repeatability of the HTML reports produced by PiGx, we identified a number of such sources of non-determinism. The Salmon aligner, for example, has a random component and does not provide a way for users to specify a seed for the pseudo-random number generators. This makes it impossible to exactly repeat an analysis and may require patching of the Salmon source code or virtualization of the random number generator facilities of the host system. Other tools are sensitive to the user’s locale settings and may generate output in non-deterministic order. We were also surprised to find that an increasingly large number of tools rely on a connection to the Internet, either directly or indirectly through dependent packages. This can be a great source of non-determinism if the experimental setup does not take the volatile nature of networked resources into account. Another important obstacle to reproducibility is the large kernel binary at runtime. Although the GNU C library provides a unified interface for all applications to use, the features that are actually implemented by the kernel at runtime may differ vastly. For example, the variant of Linux provided by Red Hat for their series 6 of operating systems reports its version as the obsolete and unsupported 2.6.32, but it contains many backported features from much newer kernel versions. Although this is usually not a problem, the kernel version and the implemented features should be taken into account. Our use of version 2.26 of the GNU C library, for example, necessitates either the use of Linux version 3.10 or higher, or a patched C library.

The use of a principled, declarative mechanism to managing software environments is a fundamental component in a holistic approach to reproducibility at all levels: repeatable builds, bit-reproducible binaries, software and data provenance, control over the configuration space, and deterministic runtime behavior. We argue that this approach can serve as a template for reproducible computational workflows.

## Acknowledgements

We are grateful to the many volunteer contributors to GNU Guix who keep improving the system.

